# Tissue micro-RNAs associated with colorectal cancer prognosis: a systematic review

**DOI:** 10.1101/701128

**Authors:** Igor Lopes dos Santos, Karlla Greick Batista Dias Penna, Megmar Aparecida dos Santos Carneiro, Larisse Silva Dalla Libera, Jéssica Enocencio Porto Ramos, Vera Aparecida Saddi

**Author notes:** Corresponding author, Address: Escola de Ciências Médicas, Farmacêutica e Biomédicas, Pontifícia Universidade Católica de Goiás, Área IV, Praça Universitária, 1440 - Setor Leste Universitário, Goiânia - GO, 74605-010, Brazil. These authors contributed equally to this work (Igor Lopes dos Santos, Vera Aparecida Saddi). These authors also contributed equally to this work (Karlla Greick Batista Dias Penna, Megmar Aparecida dos Santos Carneiro, Larisse Silva Dalla Libera, Jéssica Enocencio Porto Ramos). This research received no specific grant from any funding agency in the public, commercial, or not-for-profit sectors.

## Abstract

Colorectal cancer (CRC) is a multifactorial disease commonly diagnosed worldwide, with high mortality rates. Several studies demonstrate important associations between differential expression of micro-RNAs (miRs) and the prognosis of CRC. However, only a few systematic reviews emphasize the most relevant miRs able to contribute to the establishment of new prognostic biomarkers in CRC patients. The present study aimed to identify differentially expressed tissue miRs associated with prognostic factors in CRC patients, through a systematic review of the Literature. Using the PubMed database, Cochrane Library and Web of Science, studies published in English evaluating miRs differentially expressed in tumor tissue and significantly associated with the prognostic aspects of CRC were selected. All the included studies used RT-PCR (Taqman or SYBR Green) for miR expression analysis and the period of publication was from 2009 to 2018. A total of 115 articles accomplished the inclusion criteria and were included in the review. The studies investigated the expression of 102 different miRs associated with prognostic aspects in colorectal cancer patients. The most frequent oncogenic miRs investigated were miR-21, miR-181a, miR-182, miR-183, miR-210 and miR-224 and the hyperexpression of these miRs was associated with distant metastasis, lymph node metastasis and worse survival in patients with CRC. The most frequent tumor suppressor miRs were miR-126, miR-199b and miR-22 and the hypoexpression of these miRs was associated with distant metastasis, worse prognosis and a higher risk of disease relapse (worse disease-free survival). Specific tissue miRs are shown to be promising prognostic biomarkers in patients with CRC, given their strong association with the prognostic aspects of these tumors, however, new studies are necessary to establish the sensibility and specificity of the miRs in order to use them in clinical practice.

## Introduction

Colorectal cancer (CRC) is one of the most prevalent neoplasms in the world, being the second type of cancer more frequent in women and the third type more frequent in men. In Brazil, for the 2018-2019 biennium, it is estimated the occurrence of 36.030 new colorectal cancer cases with 17.380 cases in men and 18.980 cases in women^1^.

The survival of patients with CRC is directly associated with the pathological stage (pTNM) of the disease which is determined after microscope analysis of the sample obtained by biopsy or surgical resection of the tumor ^2, 3^. The pTNM stage is a system of stage classification proposed by the American Joint Committee on Cancer which determines the degree of tumor development according with the T, N and M categories. The T category informs about tumor growth, depth and stage of adjacent tissues invasion. The N category informs about lymph nodes metastasis and the M category describes the quantity of affected anatomical sites by distant metastasis ^3^.

On one hand, it is estimated that the survival of patients diagnosed at the first stages of the disease is up to 90%, but on the other, 30 to 45% of the patients with late diagnosis will present disease recurrence ^4, 5^. Factors such as low degree of differentiation and high levels of serum carcinoembryonic antigen (CEA) are also described as important for the increase of CRC recurrence after tumor resection ^2, 6^. However, results from recent studies have demonstrated that the analysis of micro-RNA expression in CRC can be used as a more sensible prognostic biomarker due to its potential of predicting disease recurrence, survival and therapeutic response ^7–11^.

MicroRNAs (miRs) are small non-coding RNA molecules of approximately 21 to 24 nucleotides in length that modulate gene expression by binding with the 3’ untranslated region (3’UTR) of messenger RNAs resulting in its translational repression or cleavage ^12^. In the literature, more than 2000 miRs have been already described which demonstrate an essential role in the regulation of eukaryotic cells and in the pathogenesis of several diseases, including cancer ^13, 14^.

Biogenesis of miRs begins with the activity of RNA polymerase enzyme II which transcribes the primary miRs (pri-miRs) recognized afterwards by DGCR8 protein associated with DROSHA enzyme creating a complex which allows the cleavage of the pri-miRs originating the precursor miRs (pre-miRs). The pre-miRs are carried to the cell cytoplasm by Exportin-5 and cleaved by DICER, generating double stranded miRs which will be recognized by Argonaut proteins promoting the release of one strand of the mature miR. Argonaut proteins in association with the mature miR and other proteins will constitute the RNA-Induced Silencing Complex (RISC) whose targets will be specific mRNAs. ^12, 15, 16^. Fig 1 outlines the miRs biogenesis.

**Fig 1.**
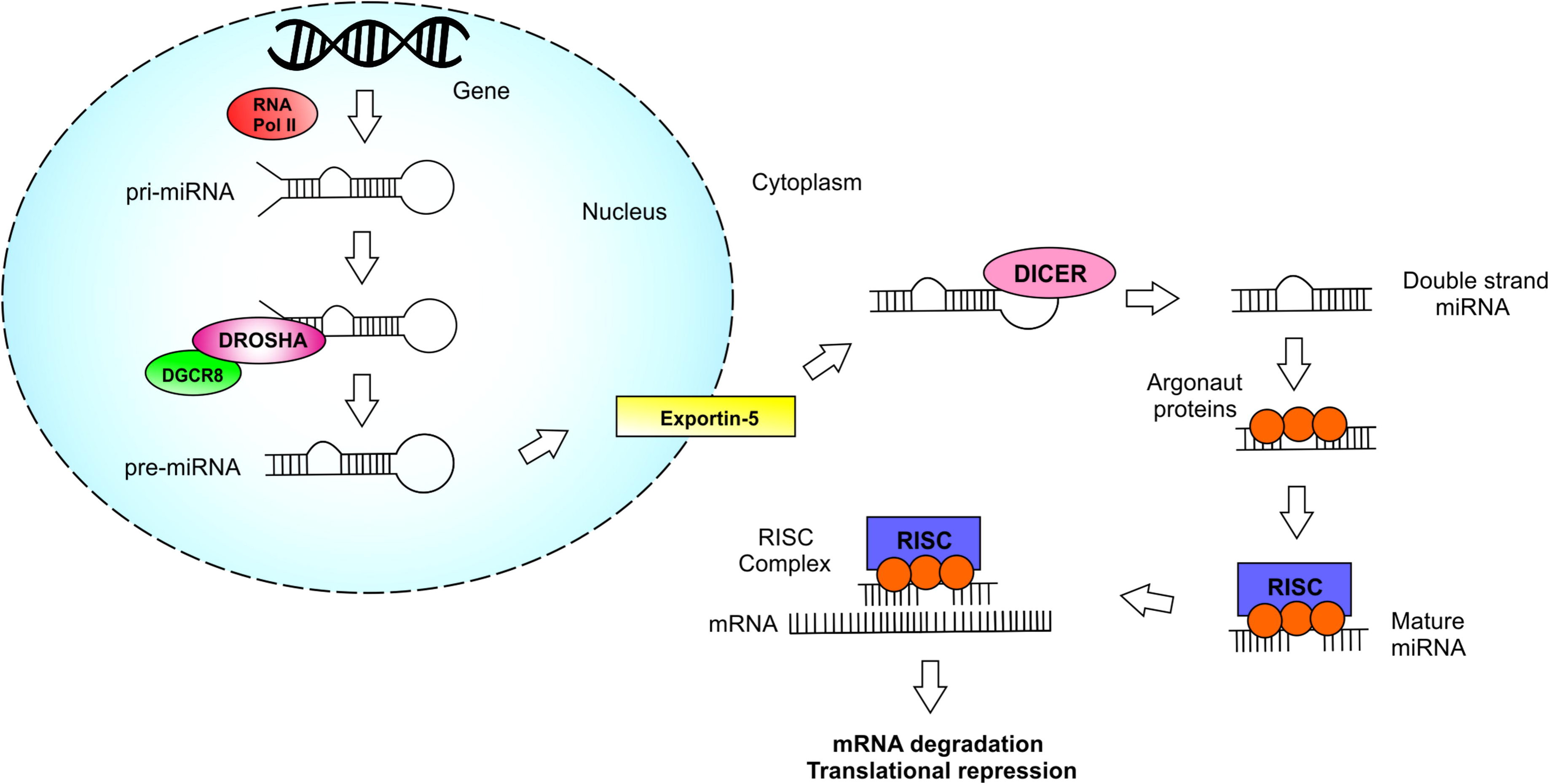
miRNAs biogenesis and proteins involved in the process.

Recent studies report that miRs regulate cellular growth and apoptosis, therefore their dysregulation could trigger carcinogenesis ^17, 18^. MiRs are classified as oncogenic or tumoral suppressor when they inhibit the expression of tumoral suppressor genes or oncogenes respectively. Oncogenic miRs are usually upregulated in the neoplastic process while tumor supressor miRs are usually downregulated in cancer ^12, 18, 19^. Consequently, studies have demonstrated by means of analysis of miR expression, that they can be valuable diagnostic and prognostic biomarkers in cancers, including CRC. ^20–23^.

Despite the recent improvements in the diagnostic methods for CRC detection, the overall survival for CRC patients is only 65% while the disease recurrence after tumor resection is approximately 45%, emphasizing the importance of describing new methods in order to guarantee a better prognosis for these patients ^4, 24^.

Several studies investigate the importance of miRs expression in CRC prognosis ^7–9, 11, 21–23^. However, there is a lack of systematic reviews in the Literature integrating and identifying the main miRs significantly associated with the prognostic aspects of CRC.

Given the relationship between the expression of miRs and the prognosis of CRC, the present study aims to highlight this association by means of a systematic review of the Literature, in order to identify the main miRs associated with each prognostic aspect of CRC. This could contribute to develop new prognostic biomarkers and could help to identify new molecular targets useful in CRC treatment.

## Methods

### Study design and search strategy

This study represents a systematic review of the Literature. For the elaboration of this project a reference review in the PubMed database, Cochrane Library and Web of Science database was performed in order to identify the relevant studies that evaluated miR expression associated with CRC prognosis. The Literature search was performed from February of 2009, year of publication of the first and less recent selected study, to December of 2018, year of publication of the most recent selected study. The following search strategy was used for the identification and selection of the articles: miRNA, colorectal cancer, prognosis. The search was performed in November 2018 and it was limited to English published articles. At the end of the search, all the results were compared and reviewed for the inclusion of relevant publications about the subject.

### Inclusion criteria

The paper selection was based on criteria related to the study classification and design in order to include the most relevant articles in this review. The relevant criteria related to the classification of the studies were: primary and descriptive studies, studies related to CRC patients and studies completely available online. The relevant criteria related to study design were: articles that evaluate CRC prognosis by overall survival and disease free survival; articles that evaluate miR expression in tumoral tissues compared to normal tissues; articles that evaluate miR expression by Real Time Polymerase Chain Reaction (RT-PCR) using Taqman and/or SYBR Green; articles that evaluate miR expression related to tumor growth/size, lymph nodes metastasis, distant metastasis and TNM stage. Publications belonging to case report, systematic reviews and meta-analysis categories were not included.

### Data Extration

The data extraction was performed by means of a collection form pre-developed by the authors in which the following data of each publication were collected: name of the first author, year of publication, micro-RNA studied, sample group, origin of cases, type of specimens analyzed, type of methodology used to evaluate the expression of micro-RNAs, target genes and association with pathological and prognostic clinical aspects.

This review followed the Preferred Reporting Items for Systematic Reviews and Meta-Analyses (PRISMA) recommendations.

### Statistical analysis and quality assessment

Due to the heterogeneity of the included studies, it was not possible to perform a weighted analysis of the results extracted from each article. Therefore, the results were analyzed qualitatively and presented narratively. A meta-analysis and the assessment of publication bias were not practically applicable for this study.

Problems with the design and execution of individual studies of healthcare interventions raise questions about the validity of their findings, and therefore, the quality of the primary studies was assessed by evaluating eight characteristics extracted from a checklist provided by the STROBE platform: description of the studying setting, description of study participants, definitions of all variables, description of the data sources and measurement tools, justification of sample size, description of statistical methods, description of results and discussion and interpretation of results. After evaluating the studies, each attribute was pointed in a range from 0 to 2 points (0 = not described, 1 = poor described, 2 = well described), with maximum quality score of 16 points in total.

### Data analysis

The data were reviewed and synthesized in semi-structured tables of Microsoft® Office Excel version 2013. The results were analyzed by descriptive statistics and qualitatively tabulated.

## Results

### Articles selection

After searching for the terms: microRNA, colorectal cancer and prognosis in all three data bases, 924 articles were found. By reading the titles, 29 duplicates were removed and 215 abstracts were selected for complete reading. After analyzing the abstracts, 134 articles which evaluate miR expression associated with CRC prognosis were selected for complete reading. After reading, 19 articles that did not achieve the proposed inclusion criteria were excluded. At the end, 115 articles were included in the systematic review. Fig. 2 outlines the study selection process.

**Fig 2.**
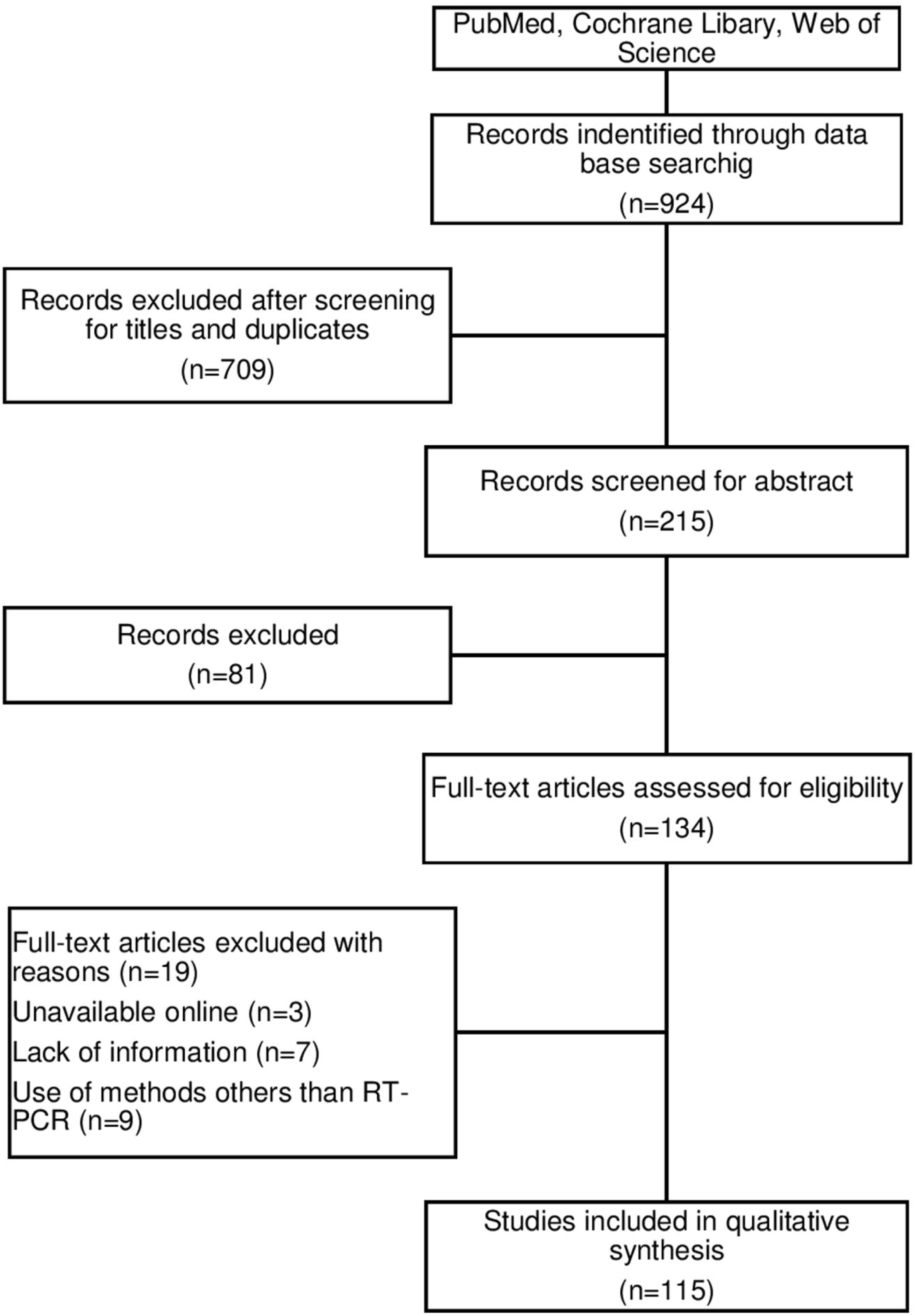
Flow diagram outlining the study selection process.

The quality scores of the studies ranged from 14 to 16 with a mean quality rating of 15.7 (S1 File).

### Characteristics of the selected studies

In total, 14.894 CRC patients were studied in the 115 included articles. The number of patients varied from 10 to 765 patients per study. The studies were performed in institutes and universities in Greece, South Korea, United States, Czech Republic, Iran, Sweden, Denmark, Japan and China with most of the research performed in the last two countries. The studies evaluated the expression of 102 miRs associated with CRC prognostics aspects. MiR-21 was the most frequent one, being analyzed in six studies ^25–30^. MiR-181a ^31–33^, miR-16 ^34–36^ and miR-133b ^37–39^ were individually investigated in three studies. MiR-126 ^8, 40^, miR-133a ^41, 42^, miR-182 ^43, 44^, miR-183 ^45, 46^, miR-199b ^47, 48^, miR-22 ^49, 50^, miR-224 ^51, 52^, miR-24-3p ^53, 54^, miR-7 ^55, 56^, were separately analyzed in two studies. Seven researches studied two miRs concomitantly ^30, 34, 38, 39, 57–59^, while the others analyzed a single miR. All studies analyzed miR expression performing RT-PCR by TaqMan and/or SYBR Green methods in which 64 studies used SYBR Green while 51 studies used Taqman. The characteristics of the selected studies in this review are available as supplemental material (S2 Table).

Studies that investigated the prognosis of CRC patients by means of overall survival and disease-free survival demonstrated that upregulated miRs associated with worse prognosis were more frequent than downregulated miRs. In terms of pathologic clinical aspects, downregulated miRs associated with advanced TNM stage, tumor growth and lymph node metastasis were more frequent than upregulated miRs. Nonetheless, upregulated miRs were more associated with distant metastasis. The association of the miRs differentially expressed in CRC and their respective prognostic aspects are summarized in Table 1.

**Table 1.** Micro-RNAs diferentially expressed in CRC and associated prognostic aspects.

| Prognostic Aspects | Downregulated micro-RNAs | Upregulated micro-RNAs |
| --- | --- | --- |
| **Advanced TNM stage** | miR-302c<sup>67</sup>, miR-466<sup>68</sup>, miR-340<sup>69</sup>, miR-196b-5p<sup>70</sup>, miR-944<sup>71</sup>, miR-30d<sup>72</sup>, miR-22<sup>50</sup>, miR-296<sup>73</sup>, miR-384<sup>74</sup>, miR-199b<sup>48</sup>, miR-128<sup>75</sup>, miR-33b<sup>76</sup>, miR-155-5p<sup>77</sup>, miR-154<sup>78</sup>, miR-149<sup>79</sup>, miR-194<sup>80</sup>, miR-133b<sup>37</sup>, miR-15a<sup>34</sup>, miR-100<sup>81</sup>, miR-133a<sup>41</sup>, miR-335<sup>82</sup>, miR-378a-3p<sup>58</sup>, miR-378a-5p<sup>58</sup>, miR-218<sup>83</sup>, miR-16<sup>34</sup> | miR-182<sup>43,44</sup>, miR-1296<sup>84</sup>, miR-888<sup>64</sup>, miR-21<sup>29</sup>, miR-130a<sup>85</sup>, miR-7<sup>55</sup>, miR-96<sup>86</sup>, miR-20a-5p<sup>63</sup>, miR-517a<sup>87</sup>, miR-106b<sup>88</sup>, miR-320e<sup>89</sup>, miR-592<sup>90</sup>, miR-196a<sup>57</sup>, miR-196b<sup>57</sup>, miR-720<sup>91</sup>, miR-133a<sup>42</sup>, miR-630<sup>92</sup>, miR-181a<sup>31</sup>, miR-210<sup>93</sup>, miR-25<sup>94</sup>, miR-191<sup>95</sup>, miR-92a<sup>96</sup>, miR-31<sup>59</sup>, miR-552<sup>97</sup> |
| **Tumor Growth** | miR-302c<sup>67</sup>, miR-466<sup>68</sup>, let-7a-5p<sup>98</sup>, miR-132<sup>99</sup>, miR-206<sup>100</sup>, miR-33b<sup>76</sup>, miR-154<sup>78</sup>, miR-194<sup>80</sup>, miR-100<sup>81</sup>, miR-378<sup>101</sup>, miR-378a-3p<sup>58</sup>, miR-378a-5p<sup>58</sup>, miR-148b<sup>102</sup>, miR-30c<sup>103</sup> | miR-1296<sup>84</sup>, miR-193a-3p<sup>104</sup>, miR-592<sup>90</sup>, miR-720<sup>91</sup>, miR-210<sup>93</sup>, miR-182<sup>44</sup>, miR-429<sup>105</sup>, miR-125b<sup>66</sup>, miR-21<sup>26,28</sup> |
| **Lymph Node Metastasis** | miR-6852<sup>106</sup>, miR-302c<sup>67</sup>, miR-466<sup>68</sup>, miR-340<sup>69</sup>, miR-944<sup>71</sup>, miR-30d<sup>72</sup>, miR-22<sup>50</sup>, miR-296<sup>73</sup>, miR-199b<sup>47</sup>, let-7a-5p<sup>98</sup>, miR-132<sup>99</sup>, miR-497<sup>107</sup>, miR-206<sup>100</sup>, miR-128<sup>75</sup>, miR-154<sup>78</sup>, miR-149<sup>79</sup>, miR-24-3p<sup>54</sup>, miR-15a<sup>34</sup>, miR-16<sup>34,36</sup>, miR-193a-5p<sup>108</sup>, miR-100<sup>81</sup>, miR-133a<sup>41</sup>, miR-205<sup>109</sup>, miR-335<sup>82</sup>, miR-378<sup>101</sup>, miR-218<sup>83</sup>, miR-138<sup>110</sup>, miR-625<sup>111</sup>, miR-212<sup>112</sup>, miR-195<sup>113</sup>, miR-133b<sup>37</sup>, miR-30c<sup>103</sup> | miR-1296<sup>84</sup>, miR-200c<sup>114</sup>, miR-21<sup>26,29</sup>, miR-130a<sup>85</sup>, miR-7<sup>55</sup>, miR-96<sup>86</sup>, miR-1260b<sup>115</sup>, miR-517a<sup>87</sup>, miR-106b<sup>88</sup>, miR-320e<sup>89</sup>, miR-196a<sup>57</sup>, miR-196b<sup>57</sup>, miR-376a<sup>116</sup>, miR-720<sup>91</sup>, miR-494<sup>61</sup>, miR-630<sup>92</sup>, miR-181a<sup>31</sup>, miR-20a<sup>62</sup>, miR-210<sup>93</sup>, miR-182<sup>43,44</sup>, miR-25<sup>94</sup>, miR-191<sup>95</sup>, miR-183<sup>46</sup>, miR-199a-3p<sup>117</sup>, miR-429<sup>105</sup>, miR-92a<sup>96</sup>, miR-10b<sup>118</sup>, miR-155<sup>30</sup>, miR-552<sup>97</sup> |
| **Distant Metastasis** | miR-466<sup>68</sup>, miR-944<sup>71</sup>, miR-30d<sup>72</sup>, miR-22<sup>49,50</sup>, miR-296<sup>73</sup>, miR-140-5p<sup>119</sup>, miR-199b<sup>48</sup>, miR-155-5p<sup>77</sup>, miR-99b-5p<sup>120</sup>, miR-154<sup>78</sup>, miR-149<sup>79</sup>, miR-96-5p<sup>121</sup>, miR-133b<sup>38,39</sup>, miR-126<sup>40</sup>, miR-138<sup>110</sup>, miR-625<sup>111</sup>, miR-212<sup>112</sup> | miR-888<sup>64</sup>, miR-20a-5p<sup>63</sup>, miR-517a<sup>87</sup>, miR-320e<sup>89</sup>, miR-592<sup>90</sup>, miR-720<sup>91</sup>, miR-494<sup>61</sup>, miR-133a<sup>42</sup>, miR-210<sup>38</sup>, miR-630<sup>92</sup>, miR-17-5p<sup>60</sup>, miR-181a<sup>31</sup>, miR-25<sup>94</sup>, miR-191<sup>95</sup>, miR-183<sup>46</sup>, miR-224<sup>52</sup>, miR-199a-3p<sup>117</sup>, miR-92a<sup>96</sup>, miR-372<sup>122</sup>, miR-185<sup>39</sup>, miR-21<sup>26,28,30</sup> |
| **Worse Overall Survival** | miR-199b<sup>47,48</sup>, miR-6852<sup>106</sup>, miR-302c<sup>67</sup>, miR-466<sup>68</sup>, miR-650<sup>123</sup>, miR-196b-5p<sup>70</sup>, miR-944<sup>71</sup>, miR-30d<sup>72</sup>, miR-296<sup>73</sup>, miR-187<sup>124</sup>, let-7a-5p<sup>98</sup>, miR-132<sup>99</sup>, miR-206<sup>100</sup>, miR-148a<sup>125</sup>, miR-33b<sup>76</sup>, miR-99b-5p<sup>120</sup>, miR-154<sup>78</sup>, miR-217<sup>126</sup>, miR-149<sup>79</sup>, miR-194<sup>80</sup>, miR-7<sup>56</sup>, miR-24-3p<sup>54</sup>, miR-29b<sup>127</sup>, miR-15a<sup>34</sup>, miR-16<sup>34,36</sup>, miR-204-5p<sup>128</sup>, miR-96-5p<sup>121</sup>, miR-193a-5p<sup>108</sup>, miR-100<sup>81</sup>, miR-133a<sup>41</sup>, miR-139-5p<sup>129</sup>, miR-124-5p<sup>130</sup>, miR-335<sup>82</sup>, miR-378<sup>101</sup>, miR-126<sup>40</sup>, miR-378a-3p<sup>58</sup>, miR-378a-5p<sup>58</sup>, miR-218<sup>83</sup>, miR-138<sup>110</sup>, miR-625<sup>111</sup>, miR-212<sup>112</sup>, miR-124<sup>131</sup>, miR-22<sup>49</sup>, miR-133b<sup>39</sup>, miR-193a-3p<sup>104</sup>, miR-195<sup>113</sup>, miR-30c<sup>103</sup> | miR-181a<sup>31-33</sup>, miR-182<sup>43,44</sup>, miR-1296<sup>84</sup>, miR-888<sup>64</sup>, miR-28-5p<sup>132</sup>, miR-183<sup>45,46</sup>, miR-200c<sup>114</sup>, miR-34a<sup>133</sup>, miR-15a-5p<sup>134</sup>, miR-181c<sup>135</sup>, miR-24-3p<sup>53</sup>, miR-16<sup>35</sup>, miR-7<sup>55</sup>, miR-96<sup>86</sup>, miR-1260b<sup>115</sup>, miR-20a-5p<sup>63</sup>, miR-517a<sup>87</sup>, miR-503<sup>136</sup>, miR-106b<sup>88</sup>, miR-211<sup>137</sup>, miR-320e<sup>89</sup>, miR-592<sup>90</sup>, miR-196a<sup>57</sup>, miR-196b<sup>57</sup>, miR-376a<sup>116</sup>, miR-224<sup>51,52</sup>, miR-720<sup>91</sup>, miR-494<sup>61</sup>, miR-223<sup>138</sup>, miR-133a<sup>42</sup>, miR-133b<sup>38</sup>, miR-210<sup>38,93</sup>, miR-630<sup>92</sup>, miR-17-5p<sup>60</sup>, miR-20a<sup>62</sup>, miR-25<sup>94</sup>, miR-191<sup>95</sup>, miR-183<sup>45,46</sup>, miR-199a-3p<sup>117</sup>, miR-429<sup>105</sup>, miR-92a<sup>96</sup>, miR-372<sup>122</sup>, miR-10b<sup>118</sup>, miR-185<sup>39</sup>, miR-21<sup>25-28,30</sup>, miR-155<sup>30</sup>, miR-125b<sup>66</sup>, miR-552<sup>97</sup> |
| **Worse Disease-Free Survival** | miR-196b-5p<sup>70</sup>, miR-944<sup>71</sup>, miR-30d<sup>72</sup>, miR-22<sup>50</sup>, miR-296<sup>73</sup>, miR-199b<sup>47</sup>, miR-187<sup>124</sup>, let-7a-5p<sup>98</sup>, miR-132<sup>99</sup>, miR-29b<sup>127</sup>, miR-15a<sup>34</sup>, miR-16<sup>34</sup>, miR-34a-5p<sup>65</sup>, miR-193a-5p<sup>108</sup>, miR-138<sup>110</sup>, miR-126<sup>8</sup>, miR-212<sup>112</sup>, miR-124<sup>131</sup> | miR-888<sup>64</sup>, miR-28-5p<sup>132</sup>, miR-183<sup>45</sup>, miR-200c<sup>114</sup>, miR-34a<sup>133</sup>, miR-15a-5p<sup>134</sup>, miR-181c<sup>135</sup>, miR-24-3p<sup>53</sup>, miR-16<sup>35</sup>, miR-96<sup>86</sup>, miR-1260b<sup>115</sup>, miR-20a-5p<sup>63</sup>, miR-517a<sup>87</sup>, miR-503<sup>136</sup>, miR-106b<sup>88</sup>, miR-320e<sup>89</sup>, miR-196a<sup>57</sup>, miR-196b<sup>57</sup>, miR-224<sup>51</sup>, miR-494<sup>61</sup>, miR-181a<sup>32</sup>, miR-21<sup>25,26,30</sup>, miR-155<sup>30</sup> |

Some of the selected studies investigated the target genes of the miRs associated with the prognostic aspects of CRC. In total, 65 target genes were analyzed in which the most frequent ones were: PTEN, investigated in six studies ^29, 32, 33, 50, 60, 61^, SMAD4, investigated in three studies ^62–64^, and TP53 ^65, 66^ investigated in two studies.

### Micro-RNAs and the prognosis of colorectal cancer

#### Micro-RNAs associated with advanced TNM stage

Twenty-four studies demonstrated the downregulation of twenty-five miRs associated with advanced TNM stage: miR-302c ^67^, miR-466 ^68^, miR-340 ^69^, miR-196b-5p ^70^, miR-944 ^71^, miR-30d ^72^, miR-22 ^50^, miR-296 ^73^, miR-384 ^74^, miR-199b ^48^, miR-128 ^75^, miR-33b ^76^, miR-155-5p ^77^, miR-154 ^78^, miR-149 ^79^, miR-194 ^80^, miR-133b ^37^, miR-15a ^34^, miR-100 ^81^, miR-133a ^41^, miR-335 ^82^, miR-378a-3p ^58^, miR-378a-5p ^58^, miR-218 ^83^ and miR-16 ^34^. Two studies demonstrated the upregulation of miR-182 ^43, 44^ associated with advanced TNM stage and others 22 studies pointed 23 different upregulated miRs associated with advanced TNM stage. In detail, miR-1296 ^84^, miR-888 ^64^, miR-21 ^29^, miR-130a ^85^, miR-7 ^55^, miR-96 ^86^, miR-20a-5p ^63^, miR-517a ^87^, miR-106b ^88^, miR-320e ^89^, miR-592 ^90^, miR-196a ^57^, miR-196b ^57^, miR-720 ^91^, miR-133a ^42^, miR-630 ^92^, miR-181a ^31^, miR-210 ^93^, miR-25 ^94^, miR-191 ^95^, miR-92a ^96^, miR-31^59^ and miR-552 ^97^ have been reported.

#### Micro-RNAs associated with tumor growth

Tumor growth was associated with the downregulation of 14 miRs reported in 13 studies: miR-302c ^67^, miR-466 ^68^, let-7a-5p ^98^, miR-132 ^99^, miR-206 ^100^, miR-33b ^76^, miR-154 ^78^, miR-194 ^80^, miR-100 ^81^, miR-378 ^101^, miR-378a-3p ^58^, miR-378a-5p ^58^, miR-148b ^102^ and miR-30c ^103^. Ten studies verified the upregulation of miR-1296 ^84^, miR-193a-3p ^104^, miR-592 ^90^, miR-720 ^91^, miR-210 ^93^, miR-182 ^44^, miR-429 ^105^,miR-125b ^66^ and miR-21 ^26, 28^ associated with tumor growth as well.

#### Micro-RNAs associated with the presence of lymph node metastasis

A total of 32 studies demonstrated the association between the downregulation of 32 miRs and the presence of lymph node metastasis such as miR-6852 ^106^, miR-302c ^67^, miR-466 ^68^, miR-340 ^69^, miR-944 ^71^, miR-30d ^72^, miR-22 ^50^, miR-296 ^73^, miR-199b^47^, let-7a-5p ^98^, miR-132 ^99^, miR-497 ^107^, miR-206 ^100^, miR-128 ^75^, miR-154 ^78^, miR-149 ^79^, miR-24-3p ^54^, miR-15a ^34^, miR-16 ^34, 36^, miR-193a-5p ^108^, miR-100 ^81^, miR-133a ^41^, miR-205 ^109^, miR-335 ^82^, miR-378 ^101^, miR-218 ^83^, miR-138 ^110^, miR-625 ^111^, miR-212 ^112^, miR-195 ^113^, miR-133b ^37^ and miR-30c ^103^. Other 30 studies verified the upregulation of miR-1296 ^84^, miR-200c ^114^, miR-21 ^26, 29^, miR-130a ^85^, miR-7 ^55^, miR-96 ^86^, miR-1260b ^115^, miR-517a ^87^, miR-106b ^88^, miR-320e ^89^, miR-196a ^57^, miR-196b ^57^, miR-376a ^116^, miR-720 ^91^, miR-494 ^61^, miR-630 ^92^, miR-181a ^31^, miR-20a ^62^, miR-210 ^93^, miR-182 ^43, 44^, miR-25 ^94^, miR-191 ^95^, miR-183 ^46^, miR-199a-3p ^117^, miR-429 ^105^, miR-92a ^96^, miR-10b ^118^, miR-155 ^30^ and miR-552 ^97^ associated with the presence of lymph node metastasis.

#### Micro-RNAs associated with the presence of distant metastasis

The downregulation of miR-466 ^68^, miR-944 ^71^, miR-30d ^72^, miR-22 ^49, 50^, miR-296 ^73^, miR-140-5p ^119^, miR-199b ^48^, miR-155-5p ^77^, miR-99b-5p ^120^, miR-154 ^78^, miR-149 ^79^, miR-96-5p ^121^, miR-133b ^38, 39^, miR-126 ^40^, miR-138 ^110^, miR-625 ^111^ and miR-212 ^112^ was associated with the presence of distant metastasis in 19 studies. The upregulation of miR-888 ^64^, miR-20a-5p ^63^, miR-517a ^87^, miR-320e ^89^, miR-592 ^90^, miR-720 ^91^, miR-494 ^61^, miR-133a ^42^, miR-210 ^38^, miR-630 ^92^, miR-17-5p ^60^, miR-181a^31^, miR-25 ^94^, miR-191 ^95^, miR-183 ^46^, miR-224 ^52^, miR-199a-3p ^117^, miR-92a ^96^, miR-372 ^122^, miR-185 ^39^ and miR-21 ^26, 28, 30^ was associated with the presence of distant metastasis in 23 studies.

#### Micro-RNAs associated with worse overall survival

Among the evaluated studies, 49 demonstrated the downregulation of miR-199b ^47, 48^, miR-6852 ^106^, miR-302c ^67^, miR-466 ^68^, miR-650 ^123^, miR-196b-5p ^70^, miR-944 ^71^, miR-30d ^72^, miR-296 ^73^, miR-187 ^124^, let-7a-5p ^98^, miR-132 ^99^, miR-206 ^100^, miR-148a ^125^, miR-33b ^76^, miR-99b-5p ^120^, miR-154 ^78^, miR-217 ^126^, miR-149 ^79^, miR-194 ^80^, miR-7 ^56^, miR-24-3p ^54^, miR-29b ^127^, miR-15a ^34^, miR-16 ^34, 36^, miR-204-5p ^128^, miR-96-5p ^121^, miR-193a-5p ^108^, miR-100 ^81^, miR-133a ^41^, miR-139-5p ^129^, miR-124-5p ^130^, miR-335 ^82^, miR-378 ^101^, miR-126 ^40^, miR-378a-3p ^58^, miR-378a-5p ^58^, miR-218 ^83^, miR-138 ^110^, miR-625 ^111^, miR-212 ^112^, miR-124 ^131^, miR-22 ^49^, miR-133b ^39^, miR-193a-3p ^104^, miR-195 ^113^ and miR-30c ^103^ associated with worse prognosis (lower overall survival) of CRC patients. Fifty-six studies identified the upregulation of miR-181a ^31–33^, miR-182 ^43, 44^, miR-1296 ^84^, miR-888 ^64^, miR-28-5p ^132^, miR-183 ^45, 46^, miR-200c ^114^, miR-34a ^133^, miR-15a-5p ^134^, miR-181c ^135^, miR-24-3p ^53^, miR-16 ^35^, miR-7 ^55^, miR-96 ^86^, miR-1260b ^115^, miR-20a-5p ^63^, miR-517a ^87^, miR-503 ^136^, miR-106b ^88^, miR-211 ^137^, miR-320e ^89^, miR-592 ^90^, miR-196a ^57^, miR-196b ^57^, miR-376a ^116^, miR-224 ^51, 52^, miR-720 ^91^, miR-494 ^61^, miR-223 ^138^, miR-133a ^42^, miR-133b ^38^, miR-210 ^38, 93^, miR-630 ^92^, miR-17-5p ^60^, miR-20a ^62^, miR-25 ^94^, miR-191 ^95^, miR-183 ^45, 46^, miR-199a-3p ^117^, miR-429 ^105^, miR-92a ^96^, miR-372 ^122^, miR-10b ^118^, miR-185 ^39^, miR-21 ^25–28, 30^, miR-155 ^30^, miR-125b ^66^ and miR-552 ^97^ was associated with worse overall survival of CRC patients.

#### Micro-RNAs associated with worse disease free-survival

In total, 17 studies demonstrated the downregulation of 18 miRs associated with worse disease free-survival such as miR-196b-5p ^70^, miR-944 ^71^, miR-30d ^72^, miR-22 ^50^, miR-296 ^73^, miR-199b ^47^, miR-187 ^124^, let-7a-5p ^98^, miR-132 ^99^, miR-29b ^127^, miR-15a ^34^, miR-16 ^34^, miR-34a-5p ^65^, miR-193a-5p ^108^, miR-138 ^110^, miR-126 ^8^, miR-212 ^112^ and miR-124 ^131^. Twenty-one studies associated the upregulation of miR-888 ^64^, miR-28-5p ^132^, miR-183 ^45^, miR-200c ^114^, miR-34a ^133^, miR-15a-5p ^134^, miR-181c ^135^, miR-24-3p ^53^, miR-16 ^35^, miR-96 ^86^, miR-1260b ^115^, miR-20a-5p ^63^, miR-517a ^87^, miR-503 ^136^, miR-106b ^88^, miR-320e ^89^, miR-196a ^57^, miR-196b ^57^, miR-224 ^51^, miR-494 ^61^, miR-181a ^32^, miR-21 ^25, 26, 30^ and miR-155 ^30^ with worse disease free-survival.

### Target genes

In order to demonstrate the association among miRs and the clinical and prognostic aspects of CRC, 60 studies investigated also the putative target genes for these miRs. Sixty-five target genes were identified and, among these genes PTEN, SMAD4, P53 and KRAS were the most frequent ones. The downregulation of PTEN was associated with the upregulation of miR-181a ^32, 33^, miR-17-5p ^60^, miR-494 ^61^, miR-22 ^50^ and miR-21 ^29^. The downregulation of SMAD4 was associated with the upregulation of miR-20a ^62^, miR-20a-5p ^63^ and miR-888 ^64^. The downregulation of P53 was related to the upregulation of miR-125b ^66^ and to the downregulation of de miR-34a-5p ^65^. The upregulation of KRAS was associated with the downregulation of miR-96-5p ^121^ and miR-384 ^74^.

## Discussion

Colorectal cancer is one of the most prevalent neoplasms in the world and it is estimated that the five-year overall survival for CRC patients is only 65%^24^. Several factors associated with CRC prognosis have been already studied, however sensible and specific prognostic biomarkers capable of predicting survival and recurrence of the disease are still unknown ^139^.

Recent reviews have demonstrated strong correlations between the prognostic aspects of breast cancer ^140^ and leukemia ^141^ with miR expression, however there are few systematic reviews integrating the main miRs associated with the prognostic aspects of CRC. The present study evaluated 113 articles that investigated the differential expression of 100 tissue miRs associated with the main clinicopathological and prognostic aspects of CRC, demonstrating that such molecules may be useful as valuable prognostic biomarkers in CRC.

MiRNAs expression can be analyzed in different biological samples such as tumor tissue ^46^, blood (serum or plasma) ^142^, feces ^143^, or urine ^144^. In the present study, articles that evaluated the expression of miRs in tumor tissues were prioritized and selected. Tumor tissues can be stored in liquid nitrogen or paraffin, allowing greater stability to the miRs molecules for later analysis in retrospective studies that aim to evaluate patients’ survival. The analysis in the tumor tissues confers greater specificity to the results since circulating miRs (present in serum or plasma) may come from different anatomical sites. In addition, tumor samples allow the analysis of miRs expression in comparison to the adjacent normal tissue of the same patient providing greater reliability to the results. In addition, studies indicate that factors associated to the tumor microenvironment influence the expression of miRs making the analysis in tumors more adequate for this type of study ^145, 146^.

The expression of miRs can be evaluated by different methods such as northern blotting, immunohistochemistry, real-time PCR (RT-PCR), microarray, and others ^147^. Each method has its own particularities and limitations. In the present study, only articles that evaluated the expression of miRs in the CRC by means of RT-PCR using TaqMan or SYBR Green were selected, because these techniques often present hight sensitivity and specificity, besides allowing the comparison of a specific miR in relation to an endogenous control.

The miRs are classified as oncogenic or tumor suppressors according to their respective target genes. MiRs that inhibit the expression of tumor suppressor genes are considered oncogenic and are generally upregulated in neoplasms. The most frequent oncogenic miRs in this study were miR-21, miR-181a, miR-182, miR-183, miR-210 and miR-224 and the upregulation of most of these miRs was associated with distant metastasis, lymph node metastasis and poorer prognosis in patients with CRC.

Conversely, miRs that inhibit the expression of oncogenes are considered tumor suppressors and are generally downregulated in the carcinogenic processes. The most frequent tumor suppressor miRs in this study were miR-126, miR-199b and miR-22 and their downregulation was associated with the presence of distant metastasis, worse prognosis and a greater risk of disease relapse (worse disease-free survival). In detail, the downregulation of miR-199b and miR-22 was also associated with more advanced TNM stage and the presence of lymph node metastasis. Other miRs such as miR-133a, miR-133b, miR-16, miR-24-3p and miR-7 were also relatively frequent in the present study, however, investigations analyzing the expression of these miRs have shown controversial results. The downregulation of these five miRs was associated with a worse prognosis and with the exception of miR-7, all the other miRs were downregulated and associated with the presence of lymph node metastasis.

In addition, other 84 miRs were also identified in the present review and their differential expression was associated with distinct clinicopathological and prognostic aspects of CRC, however, these miRs were evaluated in a single study. In general, it was observed that down and upregulated miRs associated with lymph node metastasis and worse prognosis (overall survival) in CRC were the most frequently studied.

Among the miRs identified, miR-21 was the most frequent one being analyzed in six studies. In these studies, an upregulation of miR-21 in the tumor compared to the adjacent normal tissue was found in addition to a strong correlation with more aggressive CRC characteristics such as tumor size, lymph node metastasis, distant metastasis and worse prognosis. A systematic review with meta-analysis conducted by Peng et al. selected studies that analyzed the role of miR-21 as diagnostic biomarker and prognostic of CRC, and statistically demonstrated that miR-21 upregulation in the tumor was associated with worse overall survival and worse disease-free survival, as in the present review ^148^. The findings in both studies suggest the potential of miR-21 to predict survival and relapse of the disease making it promising biomarker for CRC prognosis.

Another systematic review with meta-analysis conducted by Gao et al. evaluated the prognostic value of miRs in colorectal cancer as in the present review. However, in that review, studies that analyzed the expression of miRs in different types of samples such as serum, plasma and tissue and performing other methods of detection, as immunohistochemistry, were considered, so they presented miRs not identified in the present study. Nevertheless, the findings of the review by Gao et al. and the present review demonstrate similar results regarding the analysis of miRs expression in tumor tissues associated with the prognostic aspects of CRC, that is, the upregulation of miR-181a, miR-224, miR-21 and miR-126 downregulation were also present associated with worse overall survival in CRC patients ^21^.

The molecular mechanisms by which polyps and adenomas develop to CRC are diverse. However, the major altered signal pathways responsible for this progression in CRC are: MAPK / EGFR pathway, TGFβ pathway and via P13K / AKT pathway ^149^. Coincidentally, the most frequent target genes for the miRNAs identified in the present study were also directly associated with these pathways.

The phosphatase and tensin homolog (PTEN) protein is encoded by a tumor suppressor gene responsible for the inactivation of pathways associated with cell proliferation. PTEN negatively regulates the PI3K / AKT pathway by hydrolyzing the Phosphatidylinositol-3,4,5–trisphosphate (PIP3) into Phosphatidylinositol 4,5-bisphosphate (PIP2) inactivating AKT, a protein that has antiapoptotic action. Similarly, PTEN inactivates the MAPK pathway by ERK (Extracellular Regulated Kinase) dephosphorylation, inactivating the transmission of signals through this pathway and consequently inhibiting cell proliferation. In addition, PTEN activates RAD51, a DNA-repairing protein, and CENPC (Centromere Protein C), a protein responsible for the stability of centromere and chromosomal integrity ^150^. Situations in which PTEN is downregulated result in dysregulation in the cell cycle and destabilization of the cellular DNA and can trigger carcinogenic processes like CRC. In the present study, the upregulation of miR-181a, miR-17-5p, miR494, miR-22 and miR-21 was correlated with the downregulation of PTEN gene and worse prognosis of CRC patients.

The transforming growth factor β (TGF β) is a tumor suppressor cytokinin protein known to exert important functions in the regulation of apoptosis, differentiation, migration and cell proliferation. Through the binding of TGFß to its TGFR2 receptor, numerous proteins of this pathway are phosphorylated and activated, including the SMAD4 protein. However, studies indicate that mutations or downreglation of SMAD4, can hyperactivate TGFβ pathway conferring it the potential to promote tumor growth, invasion and metastasis ^149, 151^. In the present study, upregulation of miR-20a, miR-20a-5p and miR-888 was associated with the downregulation of SMAD4 and the presence of distant metastasis, advanced TNM stage, worse overall survival and worse disease-free survival in patients with CRC.

The TP53 gene encodes the tumor suppressor protein P53 that plays important roles in cell cycle control and apoptosis. MiRs belonging to the miR-34 family can modulate the antiproliferative and pro-apoptotic activity of P53 by inhibiting oncogenes and cell cycle associated genes. Thus, downregulation of miR-34 or overexpression of oncogenic miRs that negatively regulate p53 results in a dysregulation of the cell cycle leading to a carninogenic process ^152^. In the present study, the downregulation of P53 was correlated with the upregulation of miR-125b and downregulation of miR-34a-5p, resulting in a worse prognosis for patients with CRC.

KRAS is a protein belonging to the RAS family of proteins that control the activation of several pathways related to cell proliferation, such as the MAPK pathway. MiRs considered to be tumor suppressors may act as regulators of KRAS expression. Thus, the downregulation of these miRs may increase the activation of KRAS resulting in a dysregulated cellular proliferation ^153^. The present review demonstrated that upregulation of KRAS was associated with downregulation of miR-96-5p and miR-384, which in turn was associated with an advanced TNM stage, presence of distant metastasis and worse overall survival of patients with CRC. Fig 3 outlines the differential expression of the miRs and their respective target genes.

**Fig 3.**
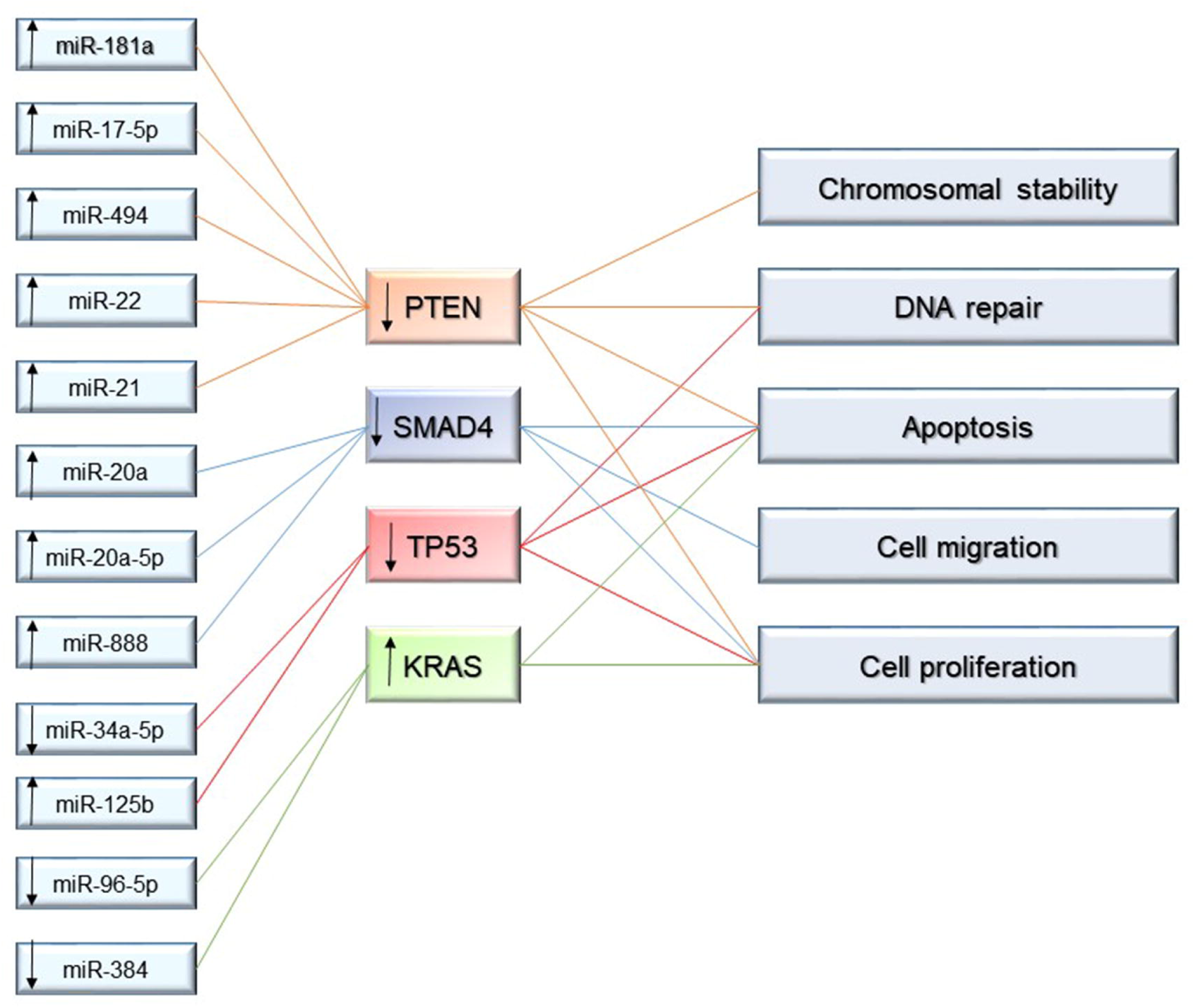
Upregulation and downregulation of miRs associated with their respective target genes and cell functions.

By analyzing the results of the present review fully, it was possible to observe the large number of miRs associated with the different prognostic aspects of CRC, making them eligible as possible prognostic markers. However, few articles have analyzed the expression of two or more miRs concomitantly. Thus, the development of studies that analyze these miRs synergistically, would allow the identification of network interactions, making them more sensitive and specific in predicting the prognosis of patients with CRC and assisting in their therapeutic behavior.

Some limitations were observed during the elaboration of the present study. The large number of studies and miRs investigated made it difficult to conduct a quantitative and qualitative analysis capable of demonstrating the most sensitive and specific differentially expressed miRs associated with each clinical-pathological and prognostic aspect of the CRC. In addition, the study was limited to articles published in English, making it harder to obtain data from studies published in other languages.

## Conclusion

After analyzing the results of the present systematic review, specific tissue miRs demonstrate to be promising CRC prognostic biomarkers due to their strong correlation with prognostic aspects of this disease, including advanced TNM stage, presence of distant metastasis and lymph node metastasis, tumor growth, prediction of worse prognosis (worse overall survival) and recurrence of the disease (worse disease-free survival).

The miR-21 was the most frequent studied miR in the present review among the miRs identified and its upregulation was associated with tumor size, lymph node metastasis, distant metastasis and worse prognosis making it a promising biomarker of CRC prognosis. However, new studies are needed to systematize and identify the most sensitive and specific miRs associated with each prognostic aspect of CRC in order to choose the most relevant ones and apply them into clinical practice.

In addition, most studies have shown strong correlations between the expression of miRs and their respective target genes, so the development of functional studies aiming to elucidate these associations becomes important as a way to help in the description of new therapeutic targets of this disease.

The vast number of miRs identified in the present study also translates the complexity and heterogeneity of CRC, emphasizing the importance of new studies that clarify the regulatory mechanisms of the expression of these miRs in this tumor, allowing a better understanding of the molecular biology of this disease.

