## Supplementary material for "Tissue micro-RNAs associated with colorectal cancer prognosis: a systematic review": S1 and S2

**Supplemental Material**

**S1. Quality assessment of the selected studies for the systematic review**

| **Ref.** | **Author / Year** | **Setting** | **Participants** | **Variables** | **Data Sources measurements** | **Study Size** | **Statistical**  **Methods** | **Results** | **Discussion** | **Total** |
| --- | --- | --- | --- | --- | --- | --- | --- | --- | --- | --- |
| (105) | Cui, 2018 | 2 | 2 | 2 | 2 | 2 | 2 | 2 | 2. | 16 |
| (83) | Tao, 2018 | 2 | 2 | 2 | 2 | 2 | 2 | 2 | 2 | 16 |
| (63) | Gao, 2017 | 2 | 2 | 2 | 2 | 2 | 2 | 2 | 2. | 16 |
| (66) | Ma, 2018 | 2 | 2 | 2 | 2 | 2 | 2 | 2 | 2 | 16 |
| (131) | Tsiakanikas, 2018 | 2 | 2 | 1 | 2 | 2 | 2 | 2 | 2 | 15 |
| (96) | Wang, 2018 | 2 | 2 | 1 | 2 | 2 | 2 | 2 | 2 | 15 |
| (44) | Chen, 2018 | 2 | 2 | 1 | 1 | 2 | 2 | 2 | 2 | 14 |
| (113) | Roh, 2018 | 2 | 2 | 1 | 2 | 2 | 2 | 2 | 2 | 15 |
| (67) | Tong, 2018 | 2 | 2 | 2 | 2 | 2 | 2 | 2 | 2 | 16 |
| (28) | Wu, 2017 | 2 | 2 | 2 | 2 | 2 | 2 | 2 | 2 | 16 |
| (84) | Chen, 2017 | 2 | 2 | 2 | 2 | 2 | 2 | 2 | 2 | 16 |
| (105) | Lin, 2017 | 2 | 2 | 2 | 2 | 2 | 2 | 2 | 2 | 16 |
| (68) | Zhang, 2017 | 2 | 2 | 2 | 2 | 2 | 2 | 2 | 2 | 16 |
| (132) | Rapti, 2017 | 2 | 2 | 1 | 2 | 2 | 2 | 2 | 2 | 15 |
| (122) | Zhou, 2017 | 2 | 2 | 2 | 1 | 2 | 2 | 2 | 2 | 15 |
| (69) | Stiegelbauer, 2017 | 2 | 1 | 2 | 1 | 2 | 2 | 2 | 2 | 14 |
| (70) | Wen, 2017 | 2 | 2 | 2 | 2 | 2 | 2 | 2 | 2 | 16 |
| (71) | Yan, 2017 | 2 | 2 | 2 | 2 | 2 | 2 | 2 | 2 | 16 |
| (49) | Xia, 2017 | 2 | 2 | 2 | 2 | 2 | 2 | 2 | 2 | 16 |
| (133) | Kontos, 2017 | 2 | 2 | 2 | 2 | 2 | 2 | 2 | 2 | 16 |
| (72) | He, 2017 | 2 | 2 | 2 | 2 | 2 | 2 | 2 | 2 | 16 |
| (134) | Yamazaki, 2017 | 2 | 2 | 1 | 2 | 2 | 2 | 2 | 2 | 15 |
| (52) | kerimis, 2017 | 2 | 2 | 1 | 2 | 2 | 2 | 2 | 2 | 15 |
| (34) | Diamantopoulos, 2017 | 2 | 2 | 1 | 2 | 2 | 2 | 2 | 2 | 15 |
| (54) | Nagano, 2016 | 2 | 2 | 1 | 2 | 2 | 2 | 2 | 2 | 16 |
| (118) | Yu, 2016 | 2 | 1 | 2 | 2 | 2 | 2 | 2 | 2 | 15 |
| (85) | Rapti, 2016 | 2 | 2 | 1 | 2 | 2 | 2 | 2 | 2 | 15 |
| (73) | Wang, 2016 | 2 | 2 | 2 | 2 | 2 | 2 | 2 | 2 | 16 |
| (46) | Cristóbal, 2017 | 2 | 1 | 2 | 2 | 2 | 2 | 2 | 2 | 15 |
| (114) | Liu , 2016 | 2 | 2 | 1 | 2 | 2 | 2 | 2 | 2 | 15 |
| (123) | Wang, 2016 | 2 | 1 | 2 | 2 | 2 | 2 | 2 | 2 | 15 |
| (62) | Cheng, 2016 | 2 | 2 | 2 | 2 | 2 | 2 | 2 | 2 | 16 |
| (97) | Liu, 2016 | 2 | 2 | 2 | 2 | 2 | 2 | 2 | 2 | 16 |
| (47) | Shen, 2016 | 2 | 2 | 2 | 2 | 2 | 2 | 2 | 2 | 16 |
| (86) | Ma, 2016 | 2 | 2 | 2 | 2 | 2 | 2 | 2 | 2 | 16 |
| (135) | Noguchi, 2016 | 2 | 1 | 2 | 2 | 1 | 2 | 2 | 2 | 14 |
| (24) | Mima, 2016 | 2 | 2 | 1 | 2 | 2 | 2 | 2 | 2 | 15 |
| (98) | Mokutani, 2016 | 2 | 2 | 2 | 2 | 2 | 2 | 2 | 2 | 16 |
| (32) | Li, 2016 | 2 | 2 | 2 | 2 | 2 | 2 | 1 | 2 | 15 |
| (102) | Bin, 2016 | 2 | 2 | 1 | 2 | 2 | 2 | 2 | 2 | 16 |
| (106) | Wang, 2016 | 2 | 2 | 2 | 2 | 2 | 2 | 2 | 2 | 16 |
| (99) | Sun, 2015 | 2 | 2 | 1 | 2 | 2 | 2 | 2 | 2 | 16 |
| (124) | Hibino, 2015 | 2 | 2 | 2 | 2 | 2 | 2 | 2 | 2 | 16 |
| (74) | Wu, 2015 | 2 | 2 | 2 | 2 | 2 | 2 | 2 | 2 | 16 |
| (75) | Liao , 2015 | 2 | 2 | 1 | 2 | 2 | 2 | 2 | 2 | 15 |
| (76) | Qu, 2015 | 2 | 2 | 1 | 2 | 2 | 2 | 2 | 2 | 15 |
| (119) | Li, 2015 | 2 | 2 | 2 | 2 | 2 | 2 | 2 | 2 | 16 |
| (87) | Zhang, 2015 | 2 | 2 | 2 | 2 | 2 | 2 | 2 | 2 | 16 |
| (136) | Sümbül, 2015 | 2 | 2 | 1 | 2 | 2 | 2 | 2 | 2 | 15 |
| (77) | Kai, 2015 | 2 | 2 | 1 | 2 | 2 | 2 | 2 | 2 | 15 |
| (88) | Perez-Carbonell, 2015 | 2 | 2 | 1 | 2 | 2 | 2 | 2 | 2 | 15 |
| (125) | Wang, 2015 | 2 | 2 | 2 | 2 | 1 | 2 | 2 | 2 | 15 |
| (89) | Liu, 2015 | 2 | 1 | 1 | 2 | 2 | 2 | 2 | 2 | 14 |
| (78) | Xu, 2015 | 2 | 2 | 2 | 2 | 1 | 2 | 2 | 2 | 15 |
| (79) | Wang, 2015 | 2 | 2 | 2 | 2 | 1 | 2 | 2 | 2 | 15 |
| (56) | Ge, 2014 | 2 | 2 | 1 | 2 | 2 | 2 | 2 | 2 | 15 |
| (55) | Suto, 2014 | 2 | 2 | 2 | 2 | 2 | 2 | 2 | 2 | 16 |
| (53) | Gao, 2015 | 2 | 2 | 1 | 2 | 2 | 2 | 2 | 2 | 15 |
| (36) | Guo, 2015 | 2 | 2 | 2 | 2 | 1 | 2 | 2 | 2 | 15 |
| (126) | Inoue, 2015 | 2 | 2 | 2 | 2 | 2 | 2 | 2 | 2 | 16 |
| (33) | Xiao, 2014 | 2 | 2 | 1 | 2 | 2 | 2 | 2 | 2 | 15 |
| (115) | Mo, 2014 | 2 | 2 | 1 | 2 | 2 | 2 | 2 | 2 | 15 |
| (25) | Fukushima, 2014 | 2 | 2 | 2 | 2 | 2 | 2 | 2 | 2 | 16 |
| (50) | Adamopoulos, 2015 | 2 | 2 | 1 | 2 | 2 | 2 | 2 | 2 | 15 |
| (64) | Gao, 2015 | 2 | 2 | 2 | 2 | 1 | 2 | 1 | 2 | 14 |
| (127) | Yin, 2014 | 2 | 2 | 2 | 2 | 2 | 2 | 2 | 2 | 16 |
| (90) | Wang, 2014 | 2 | 2 | 2 | 2 | 2 | 2 | 2 | 2 | 16 |
| (60) | Sun, 2014 | 2 | 2 | 1 | 2 | 2 | 2 | 2 | 2 | 15 |
| (137) | Li, 2014 | 2 | 2 | 1 | 2 | 2 | 1 | 2 | 2 | 14 |
| (120) | Ress, 2015 | 2 | 2 | 2 | 2 | 1 | 2 | 2 | 2 | 15 |
| (107) | Zhang, 2014 | 2 | 2 | 1 | 2 | 2 | 2 | 2 | 2 | 15 |
| (80) | Chen, 2014 | 2 | 2 | 1 | 2 | 2 | 2 | 2 | 2 | 15 |
| (40) | Wang, 2014 | 2 | 2 | 1 | 2 | 2 | 1 | 2 | 2 | 14 |
| (128) | Song, 2014 | 2 | 2 | 2 | 2 | 2 | 2 | 2 | 2 | 16 |
| (41) | Wan, 2014 | 2 | 2 | 1 | 2 | 2 | 2 | 2 | 2 | 15 |
| (129) | Jinushi, 2014 | 2 | 2 | 2 | 2 | 1 | 2 | 2 | 2 | 15 |
| (37) | Ellermeier, 2014 | 2 | 2 | 1 | 2 | 1 | 2 | 2 | 2 | 14 |
| (91) | Chu, 2014 | 2 | 2 | 1 | 2 | 2 | 2 | 2 | 2 | 15 |
| (108) | Orang, 2014 | 2 | 2 | 1 | 2 | 2 | 2 | 2 | 2 | 15 |
| (59) | Fang, 2014 | 2 | 2 | 2 | 2 | 1 | 2 | 2 | 2 | 15 |
| (81) | Sun, 2014 | 2 | 2 | 2 | 2 | 2 | 2 | 2 | 2 | 16 |
| (30) | Ji, 2014 | 2 | 2 | 2 | 2 | 2 | 2 | 2 | 2 | 16 |
| (61) | Zhang, 2014 | 2 | 2 | 2 | 2 | 2 | 2 | 2 | 2 | 16 |
| (92) | Qu, 2014 | 2 | 2 | 2 | 2 | 2 | 2 | 2 | 2 | 16 |
| (42) | Rapti, 2014 | 2 | 2 | 1 | 2 | 2 | 2 | 2 | 2 | 15 |
| (100) | Zhang, 2014 | 2 | 2 | 2 | 2 | 2 | 2 | 2 | 2 | 16 |
| (39) | Liu, 2014 | 2 | 2 | 1 | 2 | 2 | 2 | 2 | 2 | 15 |
| (57) | Li, 2014 | 2 | 2 | 1 | 2 | 1 | 2 | 2 | 2 | 14 |
| (82) | Yu, 2013 | 2 | 2 | 1 | 2 | 1 | 2 | 2 | 2 | 14 |
| (93) | Li, 2014 | 2 | 2 | 1 | 2 | 2 | 2 | 2 | 2 | 15 |
| (94) | Qin, 2014 | 2 | 2 | 2 | 2 | 2 | 2 | 2 | 2 | 16 |
| (109) | Long, 2013 | 2 | 2 | 2 | 2 | 2 | 2 | 2 | 2 | 16 |
| (45) | Zhou, 2014 | 2 | 2 | 1 | 2 | 2 | 2 | 2 | 2 | 15 |
| (35) | Qian, 2013 | 2 | 2 | 1 | 2 | 2 | 2 | 2 | 2 | 15 |
| (9) | Hansen, 2013 | 2 | 2 | 1 | 2 | 1 | 2 | 2 | 2 | 14 |
| (110) | Lou, 2013 | 2 | 2 | 1 | 2 | 2 | 2 | 2 | 2 | 15 |
| (51) | Liao, 2013 | 2 | 2 | 2 | 2 | 2 | 2 | 2 | 2 | 16 |
| (26) | Chen, 2013 | 2 | 2 | 1 | 2 | 1 | 2 | 2 | 2 | 14 |
| (27) | Toiyama, 2013 | 2 | 2 | 1 | 2 | 2 | 2 | 2 | 2 | 15 |
| (111) | Meng, 2013 | 2 | 2 | 2 | 2 | 2 | 2 | 2 | 2 | 16 |
| (43) | Liu, 2013 | 2 | 2 | 1 | 2 | 2 | 2 | 2 | 2 | 15 |
| (116) | Wan, 2013 | 2 | 2 | 1 | 2 | 2 | 2 | 2 | 2 | 15 |
| (104) | Li, 2012 | 2 | 2 | 2 | 2 | 2 | 1 | 2 | 2 | 15 |
| (31) | Nishimura, 2012 | 2 | 2 | 2 | 2 | 1 | 2 | 2 | 2 | 15 |
| (130) | Wang, 2013 | 2 | 2 | 1 | 2 | 2 | 2 | 2 | 2 | 15 |
| (95) | Zhou, 2012 | 2 | 2 | 1 | 2 | 2 | 2 | 2 | 2 | 15 |
| (48) | Zhang, 2012 | 2 | 2 | 1 | 2 | 2 | 2 | 2 | 2 | 15 |
| (121) | Yamashita, 2012 | 2 | 2 | 1 | 2 | 1 | 2 | 2 | 2 | 14 |
| (117) | Nishida, 2012 | 2 | 2 | 2 | 2 | 1 | 2 | 2 | 2 | 15 |
| (101) | Song, 2011 | 2 | 2 | 2 | 2 | 2 | 2 | 2 | 2 | 16 |
| (38) | Akçakaya, 2011 | 2 | 2 | 1 | 2 | 1 | 2 | 2 | 2 | 14 |
| (29) | Shibuya, 2011 | 2 | 2 | 2 | 2 | 2 | 2 | 2 | 2 | 16 |
| (65) | Nishida, 2011 | 2 | 2 | 2 | 2 | 1 | 2 | 2 | 2 | 15 |
| (112) | Wang, 2012 | 2 | 2 | 1 | 2 | 2 | 2 | 2 | 2 | 15 |
| (58) | Wang, 2009 | 2 | 2 | 1 | 2 | 2 | 2 | 2 | 2 | 15 |

**S2. Characteristics of the studies included in this systematic review.**

| Ref. | Author / Year | Studied micro-RNA | Target Gene | Study Population (n) | Cases Origin | Quantification Method | Results |
| --- | --- | --- | --- | --- | --- | --- | --- |
| (106) | Cui, 2018 | miR-6852 | TCF7 | CRC (n=44) and normal adjacent tissue (n=44) | Harbin, China | RT-qPCR SYBR Green | Downregulation was associated with lymph node metastasis and worse overall survival. |
| (84) | Tao, 2018 | miR-1296 | SFPQ | CRC (n= 80) and normal adjacent tissue (n= 80) | China and Japan | RT-qPCR TaqMan | Upregulation was associated with advanced TNM stage, tumor growth, lymph node metastasis and worse overall survival |
| (64) | Gao, 2017 | miR-888 | Smad4 | CRC (n= 126) and normal adjacent tissue (n= 126) | Jiangsu, China | RT-qPCR SYBR Green | Upregulation was associated with advanced TNM stage, distant metastasis, worse overall survival and worse disease free-survival. |
| (65) | Ma, 2018 | miR-302c | AP-4 | CRC (n= 90) and adjacent normal tissue (n=90) | Shaanxi, China | RT-qPCR TaqMan e SYBR Green | Downregulation was associated with advanced TNM stage, tumor depth, lymph node metastasis and worse overall survival. |
| (132) | Tsiakanikas, 2018 | miR-28-5p | / | CRC (n= 182) and adjacent normal tissue (n= 86) | Athens, Greece | RT-qPCR SYBR Green | Upregulation was associated with worse overall survival and worse disease-free survival. |
| (97) | Wang, 2018 | miR-552 | / | CRC (n= 183) and adjacent normal tissue (n= 83) | Shijiazhuan, China | RT-qPCR TaqMan | Upregulation was associated with advanced TNM stage, lymph node metastasis and worse overall survival. |
| (45) | Chen, 2018 | miR-183 | / | CRC (n=42) and adjacent normal tissue (n=42) | Tianjin, China | RT-qPCR SYBR Green | Upregulation was associated with worse overall survival and worse disease-free survival. |
| (114) | Roh, 2018 | miR-200c | / | CRC (n=109) and adjacent normal tissue (n=109) | Changwon, South Korea | RT-qPCR TaqMan | Upregulation was associated with lymph node metastasis, worse overall survival and worse disease-free survival. |
| (68) | Tong, 2018 | miR-466 | Cyclin D1, BAX e MMP-2 | CRC (n=100) and adjacent normal tissue (n=100) | Zhejiang, China | RT-qPCR SYBR Green | Upregulation was associated with advanced TNM stage, tumor growth, lymph node metastasis, distant metastasis and worse overall survival. |
| (29) | Wu, 2017 | miR-21 | PTEN | CRC (n=105) and adjacent normal tissue (n=105) | Sichuan, China | RT-qPCR SYBR Green | Upregulation was associated with advanced TNM stage and lymph node metastasis. |
| (85) | Chen, 2017 | miR-130a | Forkhead box F2 | CRC (n=53 and adjacent normal tissue (n=53) | Zhejiang, China | RT-qPCR SYBR Green | Upregulation was associated with advanced TNM stage and lymph node metastasis. |
| (104) | Lin, 2017 | miR-193a-3p | E-cadherin | CRC (n=90) and adjacent normal tissue (n=90) | Taizhou, China | RT-qPCR SYBR Green | Upregulation was associated with tumor growth and downregulation associated with worse overall survival. |
| (69) | Zhang, 2017 | miR-340 | RLIP76 | CRC (n=91) and adjacent normal tissue (n=91) | Beijing, China | RT-qPCR SYBR Green | Downregulation was associated with advanced TNM stage and lymph node metastasis. |
| (133) | Rapti, 2017 | miR-34a | / | CRC (n=133) and adjacent normal tissue (n=61) | Athens, Greece | RT-qPCR SYBR Green | Upregulation was associated with worse overall survival and worse disease-free survival. |
| (123) | Zhou, 2017 | miR-650 | AKT2/GSK3β/E-cadherin pathway | CRC patients with better prognosis (n=48) and CRC patients with worse prognosis (n=40) | Shanghai, China | RT-qPCR Taqman | Downregulation was associated with worse overall survival. |
| (70) | Stiegelbauer, 2017 | miR-196b-5p | HOXB7 e GALNT5 | Two cohorts: CRC (n=110) and CRC (n=182) | Graz, Austria e Czech Republic | RT-qPCR SYBR Green | Downregulation was associated with advanced TNM stage, worse overall survival and worse disease-free survival. |
| (71) | Wen, 2017 | miR-944 | MACC1 | CRC (n=86) and adjacent normal tissue (n=86) | Guangzhou,China | RT-qPCR TaqMan | Downregulation was associated with advanced TNM stage, lymph node metastasis, distant metastasis, worse overall survival and worse disease-free survival. |
| (72) | Yan, 2017 | miR-30d | LRH-1 | CRC (n=80) and adjacent normal tissue (n=80) | Xi'an, China | RT-qPCR SYBR Green | Downregulation was associated with advanced TNM stage, lymph node metastasis, distant metastasis, worse overall survival and worse disease-free survival. |
| (50) | Xia, 2017 | miR-22 | Sp1 ( e PTEN/AKT) | CRC (n=118) and adjacent normal tissue (n=118) | Sichuan, China | RT-qPCR TaqMan | Downregulation was associated with advanced TNM stage, lymph node metastasis, distant metastasis and worse overall survival. |
| (134) | Kontos, 2017 | miR-15a-5p | RECK | CRC (n=182) and adjacent normal tissue (n=86) | Athens, Greece | RT-qPCR SYBR Green | Upregulation was associated with worse overall survival and worse disease free-survival. |
| (73) | He, 2017 | miR-296 | S100A4 | CRC (n=90) and adjacent normal tissue (n=90) | Beijing, China | RT-qPCR TaqMan | Downregulation was associated with advanced TNM stage, lymph node metastasis, distant metastasis, worse overall survival and worse disease-free survival. |
| (135) | Yamazaki, 2017 | miR-181c | / | Two cohorts of CRC patients stage II: CRC (n= 80) and adjacent normal tissue (n=80) and CRC (n=60) and adjacent normal tissue (n=60) | Tokyo, Japan | RT-qPCR TaqMan | Upregulation was associated with worse overall survival and worse disease-free survival. |
| (53) | kerimis, 2017 | miR-24-3p | / | CRC (n=182) and adjacent normal tissue (n=86) | Athens, Greece | RT-qPCR SYBR Green | Upregulation was associated with worse overall survival and worse diasease-free survival. |
| (35) | Diamantopoulos, 2017 | miR-16 | / | CRC (n=182) and adjacent normal tissue (n=86) | Athens, Greece | RT-qPCR SYBR Green | Upregulation was associated with worse overall survival and worse diasease-free survival. |
| (55) | Nagano, 2016 | miR-7 | / | CRC (n=196) and adjacent normal tissue ((n=14) | Mie, Japan | RT-qPCR TaqMan | Upregulation was associated with advanced TNM stage, lymph node metastasis and worse overall survival. |
| (119) | Yu, 2016 | miR-140-5p | ADAMTS5, IGFBP5 | CRC (n=60) and adjacent normal tissue (n=60) | Dalian, China | RT-qPCR TaqMan | Downregulation was associated iwth tumoral stage and distant metastasis. |
| (86) | Rapti, 2016 | miR-96 | / | CRC (n=108) and adjacent normal tissue (n=54) | Athens, Greece | RT-qPCR SYBR Green | Upregulation was associated with advanced TNM stage, lymph node metastasis, worse overall survival and worse disease-free survival. |
| (74) | Wang, 2016 | miR-384 | KRAS, CDC42 | CRC (n=100) and adjacent normal tissue (n=100) | Guangdong, China | RT-qPCR SYBR Green | Downregulation was associated with advanced TNM stage. |
| (47) | Cristóbal, 2017 | miR-199b | PP2A | CRC (n=97) | Madrid, Spain | RT-qPCR TaqMan | Downregulation was associated with lymph node metastasis, worse overall survival and worse disease-free survival. |
| (115) | Liu , 2016 | miR-1260b | / | CRC (n=120) and adjacent normal tissue (n=120) | Gansu, China | RT-qPCR SYBR Green | Upregulation was associated with lymph node metastasis, worse overall survival and worse disease-free survival. |
| (124) | Wang, 2016 | miR-187 | CD276 | CRC (n=32) and adjacent normal tissue (n=32)  CRC (n=80) and adjacent normal tissue (n=80) | Shanghai, China | RT-qPCR SYBR Green | Downregulation was associated with worse overall survival and worse disease free-survival. |
| (63) | Cheng, 2016 | miR-20a-5p | Smad4 | CRC (n=544) and adjacent normal tissue (n=544) | Shanghai, China | RT-qPCR TaqMan | Upregulation was associated with advanced TNM stage, distant metastasis, worse overall survival and worse disease-free survival. |
| (98) | Liu, 2016 | let-7a-5p | HMGA2 | CRC (n=192) and adjacent normal tissue (n=192) | Taipei, Taiwan | RT-qPCR TaqMan | Downregulation was associated with tumor growth, lymph node metastasis, worse overall survival and worse diasease-free survival. |
| (48) | Shen, 2016 | miR-199b | SIRT1; CREB/KISS1 | CRC (n=60) and adjacent normal tissue (n=60) | Beijing, China | RT-qPCR SYBR Green | Downregulation associated with advanced TNM stage, distant metastasis and worse overall survival. |
| (87) | Ma, 2016 | miR-517a | FOXJ3 | CRC (n=90) and adjacent normal tissue (n=90) | Xi'an, China | RT-qPCR SYBR Green | Upregulation was associated with advanced TNM stage, lymph node metastasis, distant metastasis, worse overall survival and worse disease-free survival. |
| (136) | Noguchi, 2016 | miR-503 | CaSR | CRC (n=20), adenomas (n=20) and normal tissue (n=20) | Tsu , Japan | RT-qPCR TaqMan | Upregulation was associated with worse overall survival and worse disease-free survival. |
| (25) | Mima, 2016 | miR-21 | / | CRC (n=765) | Boston, MA, USA | RT-qPCR SYBR Green | Upregulation was associated with worse overall survival and worse disease-free survival. |
| (99) | Mokutani, 2016 | miR-132 | ANO1 | CRC (n=21 and adjacent normal tissue (n=21) | Osaka, Japan | RT-qPCR TaqMan | Downregulation was associated with tumor growth, lymph node metastasis, worse overall survival and worse disease-free survival. |
| (33) | Li, 2016 | miR-181A | PTEN/AKT | CRC (n=72) and adjacent normal tissue (n=69) | Shandong, China | RT-qPCR SYBR Green | Upregulation in hepatic metastasis patients was associated with worse overall survival. |
| (103) | Bin, 2016 | miR-30c | / | CCR (n=192) and adjacent normal tissue (n=192) | Baoding, China | RT-qPCR SYBR Green | Downregulation was associated with tumor growth, lymph node metastasis and worse overall survival |
| (107) | Wang, 2016 | miR-497 | KSR1 | CRC (n=62) and adjacent normal tissue (n=62) | Nanjing, China | RT-qPCR TaqMan | Downregulation was associated with lymph node metastasis and advanced CRC stages. |
| (100) | Sun, 2015 | miR-206 | / | CRC (n=80) and adjacent normal tissue (n=80) | Jilin, China | RT-qPCR SYBR Green | Downregulation was associated with tumor growth, lymph node metastasis and worse overall survival. |
| (125) | Hibino, 2015 | miR-148a | MMP7 | CRC (n=159) and adjacent normal tissue (n=159) | Hiroshima , Japan | RT-qPCR TaqMan | Downregulation was associated with advanced CRC stages and worse overall survival in CRC patients stage III. |
| (75) | Wu, 2015 | miR-128 | IRS1 | CRC (n=45) and adjacent normal tissue (n=45) | Changchun, China | RT-qPCR SYBR Green | Downregulation was associated with advanced TNM stage and lymph node metastasis. |
| (76) | Liao , 2015 | miR-33b | / | CRC (n=60) and adjacent normal tissue (n=60) | Shanghai, China | RT-qPCR TaqMan | Downregulation was associated with advanced TNM stage, tumor growth and worse overall survival. |
| (77) | Qu, 2015 | miR-155-5p | / | CRC (n=372) and adjacent normal tissue (n=372) | Xinjiang, China | RT-qPCR SYBR Green | Downregulation was associated with advanced TNM stage and distant metastasis. |
| (120) | Li, 2015 | miR-99b-5p | mTOR | CRC (n=56) and adjacent normal tissue (n=56) | Shanghai, China | RT-qPCR TaqMan | Downregulation was associated with distant metastasis and worse overall survival. |
| (88) | Zhang, 2015 | miR-106b | DLC1 | CRC (n=93) and adjacent normal tissue (n=93) | Sichuan, People’s Republic of China | RT-qPCR SYBR Green | Upregulation was associated with advanced TNM stage, lymph node metastasis, worse overall survival and worse disease-free survival. |
| (137) | Sümbül, 2015 | miR-211 | / | CRC (n=65), and adjacent normal tissue (n=65) and normal tissue (n=65) | Hatay, Turkey | RT-qPCR SYBR Green | Upregulation was associated with worse overall survival. |
| (78) | Kai, 2015 | miR-154 | / | CRC (n=169) and adjacent normal tissue (n=169) | Wenzhou, China | RT-qPCR TaqMan | Downregulation was associated with advanced TNM stage, tumor growth, lymph node metastasis, distant metastasis and worse overall survival. |
| (89) | Perez-Carbonell, 2015 | miR-320e | / | CRC (n=100) | Barcelona, Spain | RT-qPCR TaqMan | Up regulation was associated with advanced TNM stage, lymph node metastasis, distant metastasis, worse overall survival and worse disease-free survival. |
| (126) | Wang, 2015 | miR-217 | AEG-1 | CRC (n=50) | Beijing, P.R. China | RT-qPCR SYBR Green | Downregulation was associated with worse overall survival. |
| (90) | Liu, 2015 | miR-592 | / | CRC (n=89) and adjacent normal tissue (n=89) | Shanghai, China | RT-qPCR SYBR Green | Upregulation was associated with advanced TNM stage, tumor growth, distant metastasis and worse overall survival. |
| (79) | Xu, 2015 | miR-149 | FOXM1 | CRC (n=78) and adjacent normal tissue (n=20) | Jiangsu, PR China | RT-qPCR TaqMan | Down regulation was associated with advanced TNM stage, lymph node metastasis, distant metastasis and worse overall survival. |
| (80) | Wang, 2015 | miR-194 | MAP4K4/c- Jun/MDM2 | CRC (n=50) | Beijing, PR China | RT-qPCR SYBR Green | Downregulation was associated with advanced TNM stage, tumor growth and worse overall survival. |
| (57) | Ge, 2014 | miR-196a, miR-196b | / | CRC (n=126) and adjacent normal tissue (n=126) | Hunan, China | RT-qPCR TaqMan | Upregulation was associated with advanced TNM stage, lymph node metastasis, worse overall survival and worse disease-free survival. |
| (56) | Suto, 2014 | miR-7 | EGFR | CRC (n=105) and adjacent normal tissue (n=105) | Maebashi, Japan | RT-qPCR TaqMan | Down regulation was associated with worse overall survival. |
| (54) | Gao, 2015 | miR-24-3p | / | CRC (n=95) and adjacent normal tissue (n=95) | Shandong, China | RT-qPCR SYBR Green | Downregulation was associated with lymph node metastasis and worse overall survival. |
| (37) | Guo, 2015 | miR‑133b | CTGF | CRC (n=71) | Hunan, China | RT-qPCR SYBR Green | Downregulation was associated with lymph node metastasis and advanced TNM stage. |
| (127) | Inoue, 2015 | miR-29b | MCL1 e CDK6 | CRC (n=42) and adjacent normal tissue (n=42) | Osaka, Japan | RT-qPCR TaqMan | Downregulation was associated with worse overall survival and worse disease-free survival. |
| (34) | Xiao, 2014 | miR-15a, miR-16 | / | CRC (n=126) and adjacent normal tissue (n=126) | Hunan, China | RT-qPCR TaqMan | Downregulation was associated with advanced TNM stage, lymph node metastasis, worse overall survival and worse disease-free survival. |
| (116) | Mo, 2014 | miR-376a | / | CRC (n=53) and adjacent normal tissue (n=53) | Changchun, China | RT-qPCR SYBR Green | Upregulation was associated with lymph node metastasis and worse overall survival. |
| (26) | Fukushima, 2014 | miR-21 | / | CRC (n=306) and adjacent normal tissue (n=306) | Tokyo, Japan | RT-qPCR TaqMan | Upregulation was associated with tumor growth, lymph node metastasis, distant metastasis, worse overall survival and worse disease-free survival. |
| (51) | Adamopoulos, 2015 | miR-224 | / | CRC (n=115) and adjacent normal tissue (n=66) | Athens, Greece | RT-qPCR SYBR Green | Upregulation was associated with worse overall survival and worse disease-free survival. |
| (65) | Gao, 2015 | miR-34a-5p | p53 | CRC (n=10) and adjacent normal tissue (n=10) | Beijing, China | RT-qPCR TaqMan | Downregulation was associated with worse overall survival. |
| (128) | Yin, 2014 | miR-204-5p | RAB22A | CRC (n=272) and adjacent normal tissue (n=272) | Shanghai, China | RT-qPCR SYBR Green | Downregulation was associated with worse overall survival. |
| (91) | Wang, 2014 | miR-720 | / | CRC (n=96) and adjacent normal tissue (n=96) | Suzhou, China | RT-qPCR SYBR Green | Upregulation was associated with advanced TNM stage, tumor growth, lymph node metastasis, distant metastasis and worse overall survival. |
| (61) | Sun, 2014 | miR‑494 | PTEN | CRC (n=247) and adjacent normal tissue (n=247) | Shaanxi, china | RT-qPCR TaqMan | Upregulation was associated with lymph node metastasis, distant metastasis, worse overall survival and worse disease-free survival. |
| (138) | Li, 2014 | miR-223 | / | CRC (n=62) and adjacent normal tissue (n=62) | Jinan, China | RT-qPCR | Upregulation was associated with worse disease-free survival. |
| (121) | Ress, 2015 | MiR-96-5p | KRAS | CRC (n=80) | Graz, Austria | RT-qPCR SYBR Green | Downregulation was associated with distant metastasis and worse overall survival. |
| (108) | Zhang, 2014 | miR-193a-5p | / | CRC (n=69) and adjacent normal tissue (n=69) | Beijing, China | RT-qPCR TaqMan | Downregulation was associated with lymph node metastasis, worse overall survival and worse disease-free survival. |
| (81) | Chen, 2014 | miR-100 | / | CRC (n=138) and adjacent normal tissue (n=138) | Zaozhuang, China | RT-qPCR SYBR Green | Downregulation was associated with advanced TNM stage, tumor growth, lymph node metastasis and worse overall survival. |
| (41) | Wang, 2014 | miR-133a | / | CRC (n=169) and adjacent normal tissue (n=169) | Shandong, China | RT-qPCR | Downregulation was associated with advanced TNM stage, lymph node metastasis and worse overall survival. |
| (129) | Song, 2014 | miR-139-5p | AMFR and NOTCH1 | CRC (n=158) and adjacent normal tissue (n=158) | Shanghai, China | RT-qPCR SYBR Green | Downregulation was associated with worse overall survival. |
| (42) | Wan, 2014 | miR-133a | / | CRC (n=125) and adjacent normal tissue (n=125) | Hong Kong, China | RT-qPCR SYBR Green | Upregulation was associated with advanced TNM stage, distant metastasis and worse overall survival. |
| (130) | Jinushi, 2014 | miR-124-5p | SMC4 | CRC (n=71) | Sapporo, Japan | RT-qPCR SYBR Green e TaqMan | Downregulation was associated with worse overall survival. |
| (38) | Ellermeier, 2014 | miR-133b e miR-210 | / | CRC (n=19 and adjacent normal tissue (n=19) | Providence, RI, U.S.A | RT-qPCR TaqMan | Upregulation of both was associated with worse overall survival. Upregulation of miR-210 and downregulation of miR-133b were associated with distant metastasis. |
| (92) | Chu, 2014 | miR-630 | / | CRC (n=206) and adjacent normal tissue (n=206) | Xi’na, China | RT-qPCR TaqMan | Upregulation was associated with advanced TNM stage, lymph node metastasis, distant metastasis and worse overall survival. |
| (109) | Orang, 2014 | miR-205 | / | CRC (n=40) and adjacent normal tissue (n=40) | Tabriz, Iran | RT-qPCR SYBR Green | Downregulation was associated with lymph node metastasis. |
| (60) | Fang, 2014 | miR-17-5p | PTEN | CRC (n=295) | Guangzhou, China | RT-qPCR SYBR Green | Upregulation was associated with distant metastasis and worse overall survival. |
| (82) | Sun, 2014 | miR-335 | ZEB2 | CRC (n=80) and adjacent normal tissue (n=80) | Xi’na, China | RT-qPCR SYBR Green | Downregulation was associated with advanced TNM stage, lymph node metastasis and worse overall survival. |
| (31) | Ji, 2014 | miR-181a | WIF-1 | CRC (n=137) and adjacent normal tissue (n=137) | Beijing, China | RT-qPCR TaqMan | Upregulation was associated with advanced TNM stage, lymph node metastasis, distant metastasis and worse overall survival. |
| (62) | Zhang, 2014 | miR‑20a | SMAD4 | CRC (n=86) and adjacent normal tissue (n=86) | Nanchong, China | RT-qPCR SYBR Green | Upregulation was associated with lymph node metastasis and worse overall survival. |
| (93) | Qu, 2014 | miR-210 | VMP1 | CRC (n=193) and adjacent normal tissue (n=193) | Shandong, China | RT-qPCR SYBR Green | Upregulation was associated with advanced TNM stage, tumor growth, lymph node metastasis and worse overall survival. |
| (43) | Rapti, 2014 | miR-182 | / | CRC (n=116) and adjacent normal tissue (n=60) | Athens, Greece | RT-qPCR SYBR Green | Upregulation was associated with advanced TNM stage, lymph node metastasis and worse overall survival. |
| (101) | Zhang, 2014 | miR-378 | Vimentin | CRC (n=84) and adjacent normal tissue (n=84) | Sichuan, China | RT-qPCR SYBR Green | Downregulation was associated with tumor growth, lymph node metastasis and worse overall survival. |
| (40) | Liu, 2014 | miR-126 | / | CRC (n=92) and adjacent normal tissue (n=92) | Zhanjiang, China | RT-qPCR SYBR Green | Downregulation was associated with distant metastasis and worse overall survival. |
| (58) | Li, 2014 | miR-378a-3p, miR-378a-5p | / | Two cohorts: CRC (n=96) and adjacent normal tissue (n=96);  CRC (n=12) and adjacent normal tissue (n=12) | Guangzhou, China | RT-qPCR SYBR Green | Downregulation was associated with advanced TNM stage, tumor growth and worse overall survival. |
| (83) | Yu, 2013 | miR-218 | / | CRC (n=26) and adjacent normal tissue (n=26) | Jiangsu, China | RT-qPCR SYBR Green | Downregulation was associated with advanced TNM stage, lymph node metastasis and worse overall survival. |
| (94) | Li, 2014 | miR-25 | / | CRC (n=186) and adjacent normal tissue (n=186) | Shaanxi, China | RT-qPCR TaqMan | Upregulation was associated with advanced TNM stage, lymph node metastasis, distant metastasis and worse overall survival. |
| (95) | Qin, 2014 | miR-191 | TIMP3 | CRC (n=136) | Xinxiang, China | RT-qPCR TaqMan | Upregulation was associated with advanced TNM stage, lymph node metastasis, distant metastasis and worse overall survival. |
| (110) | Long, 2013 | miR-138 | TWIST2 | CRC (n=187) and adjacent normal tissue (n=187) | Hunan,China | RT-qPCR TaqMan | Downregulation was associated with lymph node metastasis, distant metastasis, worse overall survival and worse disease-free survival. |
| (46) | Zhou, 2014 | miR-183 | / | CRC (n=94) and adjacent normal tissue (n=94) | Sichuan, China | RT-qPCR SYBR Green | Upregulation was associated with lymph node metastasis, distant metastasis and worse overall survival. |
| (36) | Qian, 2013 | miR-16 | / | CRC (n=143) and adjacent normal tissue (n=18) | Jiangsu, China | RT-qPCR TaqMan | Downregulation was associated with advanced TNM stage, lymph node metastasis and worse overall survival. |
| (8) | Hansen, 2013 | miR-126 | / | CRC (n=126) | Odense, Denmark/ Lund, Sweden | RT-qPCR SYBR Green | Downregulation was associated with worse overall survival. |
| (111) | Lou, 2013 | miR-625 | / | CRC (n=96) and adjacent normal tissue (n=96) | Shanghai, China | RT-qPCR SYBR Green | Downregulation was associated with lymph node metastasis, distant metastasis and worse overall survival. |
| (52) | Liao, 2013 | miR-224 | PHLPP1, PHLPP2 | CRC (n=43) and adjacent normal tissue (n=43) | Guangzhou, China | RT-qPCR TaqMan | Upregulation was associated with distant metastasis and worse overall survival. |
| (27) | Chen, 2013 | miR-21 | / | CRC (n=195) | Changhua, Taiwan | RT-qPCR TaqMan | Upregulation was associated with worse overall survival. |
| (28) | Toiyama, 2013 | miR-21 | / | CRC (n=166) and adjacent normal tissue (n=166) | Japan | RT-qPCR TaqMan | Upregulation was associated with tumor growth, distant metastasis and worse overall survival. |
| (110) | Meng, 2013 | miR-212 | MnSOD | CRC (n=30) and adjacent normal tissue (n=30) | Guangzhou, China | RT-qPCR TaqMan | Downregulation was associated with lymph node metastasis, distant metastasis, worse overall survival and worse disease-free survival. |
| (44) | Liu, 2013 | miR-182 | / | CRC (n=152) and adjacent normal tissue (n=152) | Jinan, China | RT-qPCR SYBR Green | Upregulation was associated with advanced TNM stage, tumor growth, lymph node metastasis and worse overall survival. |
| (117) | Wan, 2013 | miR-199a-3p | / | CRC (n=92) and adjacent normal tissue (n=92) | Shanghai, China | RT-qPCR SYBR Green | Upregulation was associated with lymph node metastasis, distant metastasis and worse overall survival. |
| (105) | Li, 2012 | miR-429 | SOX2 | CRC (n=107) and adjacent normal tissue (n=107) | Shandong, China | RT-qPCR | Upregulation was associated with tumor growth, lymph node metastasis and worse overall survival. |
| (32) | Nishimura, 2012 | miR-181a | PTEN | CRC (n=162) | Beppu, Japan | RT-qPCR TaqMan | Upregulation was associated with worse overall survival and worse disease-free survival. |
| (131) | Wang, 2013 | miR-124 | / | CRC (n=96) and adjacent normal tissue (n=96) | Chengdu City, China | RT-qPCR TaqMan | Downregulation was associated with worse overall survival and worse disease-free survival. |
| (96) | Zhou, 2012 | miR-92a | / | CRC (n=82) and adjacent normal tissue (n=82) | Nanchong, China | RT-qPCR SYBR Green | Upregulation was associated with advanced TNM stage, lymph node metastasis, distant metastasis and worse overall survival. |
| (49) | Zhang, 2012 | miR-22 | / | CRC (n=86) and adjacent normal tissue (n=86) | Nanchong, China | RT-qPCR SYBR Green | Downregulation was associated with distant metastasis and worse overall survival. |
| (122) | Yamashita, 2012 | miR-372 | / | CRC (n=144) | Beppu, Japan | RT-qPCR TaqMan | Upregulation was associated with distant metastasis and worse overall survival. |
| (118) | Nishida, 2012 | miR-10b | BIM | CRC (n=88) | Osaka, Japan | RT-qPCR TaqMan | Upregulation was associated with lymph node metastasis and worse overall survival. |
| (102) | Song, 2011 | miR-148b | CCK2R | CRC (n=96) and adjacent normal tissue (n=96) | Shenyang City, China | RT-qPCR SYBR Green | Downregulation was associated with tumor growth. |
| (39) | Akçakaya, 2011 | miR-185, miR-133b | / | CRC (n=50) | Stockholm, Sweden | RT-qPCR SYBR Green e TaqMan | Upregulation of miR-185 and downregulation of miR-133b were associated with metastasis and worse overall survival. |
| (30) | Shibuya, 2011 | miR-21 e miR-155 | PDCD4 and TP53INP1 | CRC (n=156) and adjacent normal tissue (n=156) | Tokyo, Japan | RT-qPCR TaqMan | Upregulation of miR-21 was associated with distant metastasis, worse overall survival and worse disease-free survival. Upregulation of miR-155 as associated with lymph node metastasis, worse overall survival and worse disease-free survival. |
| (66) | Nishida, 2011 | miR-125b | p53 e p21 | CRC (n=89) | Osaka, Japan | RT-qPCR TaqMan | Upregulation was associated with tumor growth and worse overall survival. |
| (113) | Wang, 2012 | miR-195 | / | CRC (n=85) and adjacent normal tissue (n=85) | Nanjing, China | RT-qPCR SYBR Green | Downregulation was associated with lymph node metastasis and worse overall survival. |
| (59) | Wang, 2009 | miR-31, miR-143, miR-145 | / | CRC (n=98) and adjacent normal tissue (n=98) | Chengdu, China | RT-qPCR TaqMan | Upregulation of miR-31 was associated with advanced TNM stage. |

miR: micro-RNA. CRC: colorectal cancer. RT-qPCR: real-time polymerase chain reaction
